# Defining a landscape of molecular phenotypes using a simple single sample scoring method

**DOI:** 10.1101/231217

**Authors:** Momeneh Foroutan, Dharmesh D. Bhuva, Kristy Horan, Ruqian Lyu, Joseph Cursons, Melissa J. Davis

## Abstract

**Background:** Gene set scoring provides a useful approach for quantifying concordance between sample transcriptomes and selected molecular signatures. Most methods use information from all samples to score an individual sample, leading to unstable scores in small data sets and introducing biases from sample composition across a data set (e.g. varying numbers of samples for different cancer subtypes). To address these issues we have developed a truly single sample scoring method, and associated *R/Bioconductor* package *singscore*.

**Results:** We have developed a rank-based single sample scoring method, implemented as a Bioconductor package. We use multiple cancer data sets to compare it against widely-used scoring methods, including GSVA, z-scores, PLAGE, and ssGSEA. Our approach does not depend upon background samples and thus the scores are stable regardless of the composition and number of samples in the gene expression data set. In contrast, scores obtained by GSVA, *z*-score, PLAGE and ssGSEA can be unstable when less data are available (*n_samples_* < 25). We show that the computational time for *singscore* is faster than current implementations of GSVA and ssGSEA, and is comparable with that of z-score and PLAGE. The *singscore* package also produces visualisations and interactive plots that enable exploration of molecular phenotypes.

**Conclusions:** The single sample scoring method described here is independent of sample composition in gene expression data and thus it provides stable scores that are less likely to be influenced by unwanted variation across samples. These scores can be used for dimensional reduction of transcriptomic data and the phenotypic landscapes obtained by scoring samples against multiple molecular signatures may provide insights for sample stratification.

## Background

Several approaches have been developed to score individual samples against molecular signatures (or gene sets), including: ssGSEA (single sample gene set enrichment analysis) [2], GSVA (gene set variation analysis) [3], PLAGE (pathway level analysis of gene expression) [4] and combining z-scores [5].

Hänzelmann et al. (2013) implemented all four of these methods within the *R/Bioconductor* package GSVA and performed a detailed comparison. It should be noted that GSVA, PLAGE and *z*-scores use data from all samples in the very first step to estimate gene distributions; GSVA performs kernel density estimation of the expression profile for each gene across all samples, while PLAGE and z-scores apply standardisation methods. Although ssGSEA normalises the scores across samples, this is the final step and it can be disabled. Some methods also make assumptions about the data which may be unsuitable in certain cases, for instance, PLAGE and combined *z*-scores are parametric methods that assume normality of expression profiles, while the combined *z*-scores method additionally makes an independence assumption for genes in a gene set [3].

Here, we introduce a rank-based single sample scoring method, *singscore*. Using breast cancer data and several gene expression signatures we compare our approach to the above methods. The *singscore* method is simple, making the scores directly interpretable (as a normalised mean percentile rank), and our comparisons show that it is not only fast, but it also produces stable and reproducible scores regardless of the composition and number of samples within the data. Due to the independence of samples in computing scores, when within-sample effects such as RNA-seq gene length bias [6, 7], GC content bias [8], and other technical artefacts are properly corrected [9-11], we believe this method may be more robust to unwanted variation that exists across samples. Furthermore, we include examples from real cancer data to show visualisation options and applications of the simple scoring method in molecular phenotyping and a clinical context.

## Methods

### The singscore method

For a sample transcriptome which has been corrected for technical within-sample bias (i.e. RPKM, TPM, or RSEM data for RNA-seq), genes are ranked by increasing mRNA abundance. Given an expression signature with expected up regulated genes the mean rank of this gene set in the ordered data is calculated; if the gene set also has a down regulated component, the ranks are reversed before the mean rank of the down-set is calculated. Mean ranks are separately normalised (relative to theoretical minimum and maximum scores), and then summed (if an expected downregulated set is present), before centering on zero. A sample with a high score can be interpreted as having a transcriptome which is concordant to the specified signature, and scores reflect the relative mean percentile rank of the target gene sets within each sample.

Mathematically, the scores are defined as:

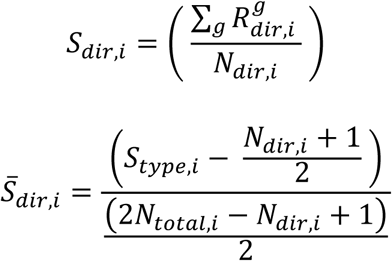

Where:

- *dir* is the gene set direction (*i.e*. expected up- or down-regulated genes);
- *S_dir,i_* is the score for sample *i* against the directed gene set;
- 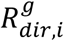 is the rank of gene *g* in the directed gene set (decreasing abundance for expected upregulated genes and decreasing abundance for expected downregulated genes);
- *N_dir,i_* is number of genes in the expected up- or down-regulated gene set that are observed within the data (*i.e*. missing genes are excluded);
- 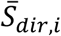 is the normalised score for sample *i* against those genes, and;
- *N_total,i_* is the number of total genes in sample *i*.

As noted above, the up- and down-scores for each sample are normalized against the theoretical minimum and maximum values calculated under the assumption of non-overlapping ranks (determined from the sum of arithmetic series), summed, and then centered on zero. A static centering is performed in a score-independent manner such that sample independence is maintained. If required, the significance of a score can be evaluated using a permutation test, where scores are computed per sample for random gene sets of the same size as the set of interest to form a null distribution (Fig. S1 *in Additional File* 1). The approach is similar to a Wilcoxon rank sum test if the gene-set consists of either one of up-regulated or down-regulated genes.

### Implementation of singscore

All statistical analyses were performed using *R* (v3.3 and greater) and Bioconductor (v3.4 and greater). We have produced a `singscore` package implementing this method, and included several visualisation functions that produce both static and interactive (*.html*) plots using `ggplot2` [12] and `plotly` [13] respectively.

### Other scoring methods

The *R/Bioconductor* package `GSVA` was used to evaluate the performance of the GSVA, ssGSEA, z-score and PLAGE methods [14]. We have slightly modified this approach to account for bidirectional signatures where both expected up- and down-regulated gene sets were available, with a method previously described by Foroutan et al. [1]. Additionally, we include ssGSEA_!Norm_ by removing the (final) normalisation step of the ssGSEA method, allowing us to test the performance in data sets with smaller sample sizes.

### Data

In this study, we used The Cancer Genome Atlas (TCGA) breast cancer [15] RNA-seq data (RSEM normalised) and microarray data (RMA normalised from Agilent4502A_07_03 microarray platform), the Cancer Cell Line Encyclopaedia (CCLE) [16] breast cancer cell line RNA-seq data (TPM normalised), the raw fastq files of the breast cancer cell lines from A Daemen, OL Griffith, LM Heiser, NJ Wang, OM Enache, Z Sanborn, F Pepin, S Durinck, JE Korkola, M Griffith, et al. [17] (re-calculated as RPKM; *see Data processing below*), and the integrated cell line TGFβ-EMT data from M Foroutan, J Cursons, S Hediyeh-Zadeh, EW Thompson and MJ Davis [1].

### Data processing

The SRA files from Daemen *et al*. were obtained on July 2016 (GSE), and converted to fastq files using the `fastq-dump` function in the SRA toolkit [21]. Reads were aligned to human genome hg19 using the `Rsubread` package [22] in *R/Bioconductor*, and count level data were obtained using the `featureCount` function with default parameters. The `edgeR` package [23] was used to calculate RPKM values. For all other data sets, we used processed data available online (see Table 1).

**Table 1.** List of data sets used in the current study.

| Data | Source | Date downloaded | Reference |
| --- | --- | --- | --- |
| TCGA RNA-seq | The UCSC Cancer Genomics Browser [18] | February 2016 | PMID: 23000897 |
| TCGA microarray | The UCSC Cancer Genomics Browser [18] | October 2015 | PMID: 23000897 |
| CCLE RNA-seq | Cancer Cell Line Data Repository [19] | April 2017 | PMID: 22460905 |
| Daemen et al RNA-seq | Gene Expression Omnibus [20] ID: GSE48213 | August 2016 | PMID: 24176112 |
| TGF $\beta$ -EMT | [doi: 10.4225/49/5a2a11fa43fe3](https://doi.org/10.4225/49/5a2a11fa43fe3) | - | PMID: 28119430 |

### Comparing the stability of single scores obtained by scoring methods

Method stability was examined using 500 of the overlapping samples between the TCGA breast cancer RNA-seq and microarray data (Sample IDs in Suppl. Table 1). Sub-sampling was then performed to vary the number of samples and genes present for each evaluation. To examine sample size effects upon a given sample, *s_i_*, two data sets were created by sampling from both the RNA-seq and microarray data to select the sample *s_i_* and *n* – 1 other random samples. The score for sample *s_i_* was then computed using all listed methods, and this process was repeated across all 500 samples at a given sample size, such that there are 500 matched scores in total from each of the microarray data and RNA-seq data. The Spearman’s rank correlation coefficient and the concordance index were then calculated between sample scores from the microarray and the RNA-seq data. We note that for some methods, sampling data in such a manner can modify the background samples for a sample of interest, reflecting the effect of overall sample composition on the final scores. A similar analysis was performed by varying the number of genes in the sub-sampled data, however, note that when sampling genes from the original data, all genes present in the signature of interest were retained.

We performed this analysis with undirected epithelial and mesenchymal gene sets [24], and a directed TGFβ-EMT signature [1], using test combinations of sample numbers *n* = (2, 5,25, 50, 500) and gene numbers *N_G_* = (500,1000, 5000,10000, *ALLGENES*). The complete set of permutations was repeated 20 times to ensure accuracy of estimates and to estimate the error margins of measurements.

### Comparing the computation time for scoring methods

To compare the computational time of each scoring method, we randomly selected 1000 gene sets from MSigDB [25, 26] and scored the TCGA breast cancer RNA-seq data using all methods listed above. The analysis was repeated 10 times to improve coverage of signatures on MSigDB and allow error estimates. This comparison was performed on UNIX machine (Intel(R) Xeon ^®^ CPU E5-2690 v3 @ 2.60GHz) using only one core without code parallelisation.

## Results

### Technical considerations for simple scoring method

#### Simple scores are highly stable compared to other scoring approaches

The performance of *singscore* was compared to GSVA, z-score, PLAGE, ssGSEA, and a modified ssGSEA without normalization (ssGSEA_!Norm_), using both the TCGA breast cancer microarray and RNA-seq data. Overlapping samples between the two platforms (*n_samples_* = 500) were scored using three gene signatures, epithelial (Epi), mesenchymal (Mes) and TGFβ-induced EMT (TGFβ-EMT) signatures [24] while the number of samples and genes in the data were modified (*as described in the Methods section*). The Spearman’s correlations and concordance index [27] between sample scores from the two platforms were calculated. Our results show a high stability for *singscore* and ssGSEA modified to run without normalisaton (ssGSEA_!Norm_) compared to the other methods when varying the sample number and number of genes in the data set (Figure 1, Suppl. Figures 2 and 3) reflecting sample composition effects. While all methods performed well for large data sets, PLAGE had the worst performance with sub-sampled data, while GSVA, z-score and ssGSEA show a reduced stability compared to *singscore* and ssGSEA_!Norm_ in data sets with small sample sizes (*n_Samples_* < 25). This demonstrates that *singscore* may be particularly useful in cases where data sets are relatively small, or where there may be a heterogeneous sample composition (*i.e*. samples across different cancer subtypes with unbalanced frequencies). Although PLAGE appeared to perform poorly for signatures with both expected up- and down-regulated gene sets, as discussed below, the underlying metric was not designed to account for such directionality.

**Figure 1.**
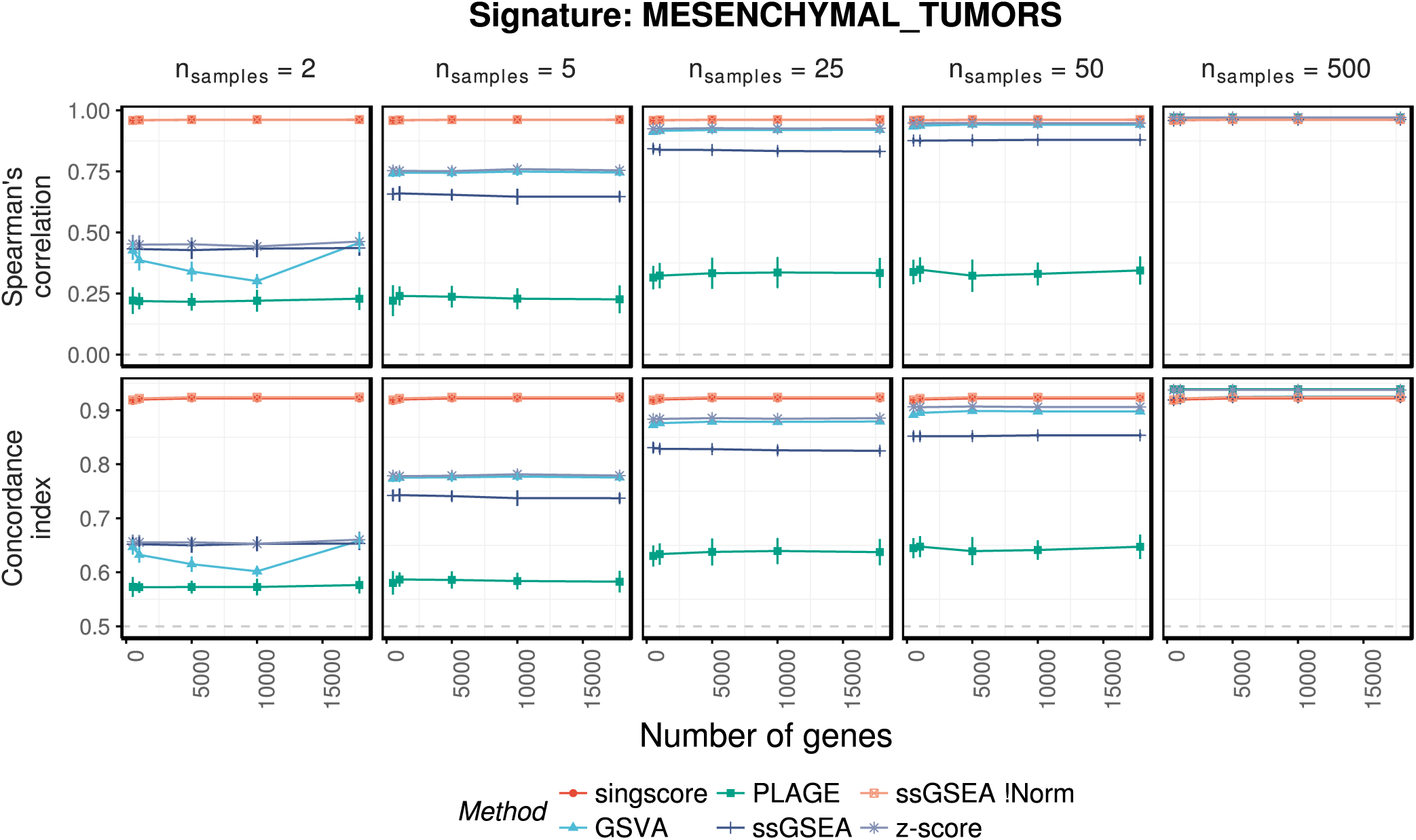
Comparing the stability of scoring methods to changes in number of samples and genes in the transcriptomic data sets. For both Spearman’s correlation coefficients and concordance index (C-Index), a method with a higher value indicates better performance with 0 and 0.5 indicating poor performance respectively. Similar results are observed when other signatures were used for scoring (Suppl. Figures 2 and 3).

An important factor for computational tools is run time, and we note that ssGSEA_!Norm_ and ssGSEA have much longer compute times than all other methods when tested with random signatures from MSigDB [25,26] (Method section, Suppl. Figure 1), whereas our approach is very fast and comparable with PLAGE and z-score.

#### Scores obtained by *singscore are* not influenced by (unwanted) variation across samples

An important consideration with transcriptomic data is the presence of (unwanted) variation across or within some samples (*i.e*. batch effects); for methods that use all the samples within a dataset, the score of an individual sample can thus be affected by artefacts within other samples. Further, for cancer data sets, the scores of each sample will be affected by the relative composition of the overall study (*i.e*. they could be affected by including/excluding different subtypes). Our comparison between scores derived from the microarray (*n_Genes_* = 17814) and RNA-seq (*n_Genes_* = 14202) expression data for varying numbers of samples (*n_Samples_* = 2-500, Figure 1) shows changes in the scores of an individual sample with different sample backgrounds, when calculated by PLAGE, GSVA, z-score, and ssGSEA methods, particularly with smaller data sets; while the *singscore* and ssGSEA_!Norm_ metrics are more robust to changes in sample composition and size, due to the independence of scores between samples.

We also compared independent transcriptomic data for cell lines collected from two independent studies [16, 17], and calculated both the epithelial and mesenchymal scores for the cell lines. There were 32 cell lines present in each dataset, which enabled us to explore the consistency of EMT scoring in independent datasets. The scores are largely consistent (Figure 2A), despite differences in computational pipelines, gene expression metrics and experimental protocol of the two datasets. For the few cell lines that show substantial variation in scores we cannot exclude the possibility that biological variation between the independently cultured cells might underpin the observed differences. We note that all the cell lines with larger variations in mesenchymal scores are from luminal sub-groups with consistently high epithelial scores across the two datasets, while cell lines with the highest variability in epithelial scores are from the claudin-low sub-groups which also show consistently high mesenchymal scores (Figure 2A). The variation in epithelial and mesenchymal scores for three representative cell lines (HCC1428, HCC202, and MDAMB231) is shown in Figure 2B.

**Figure 2.**
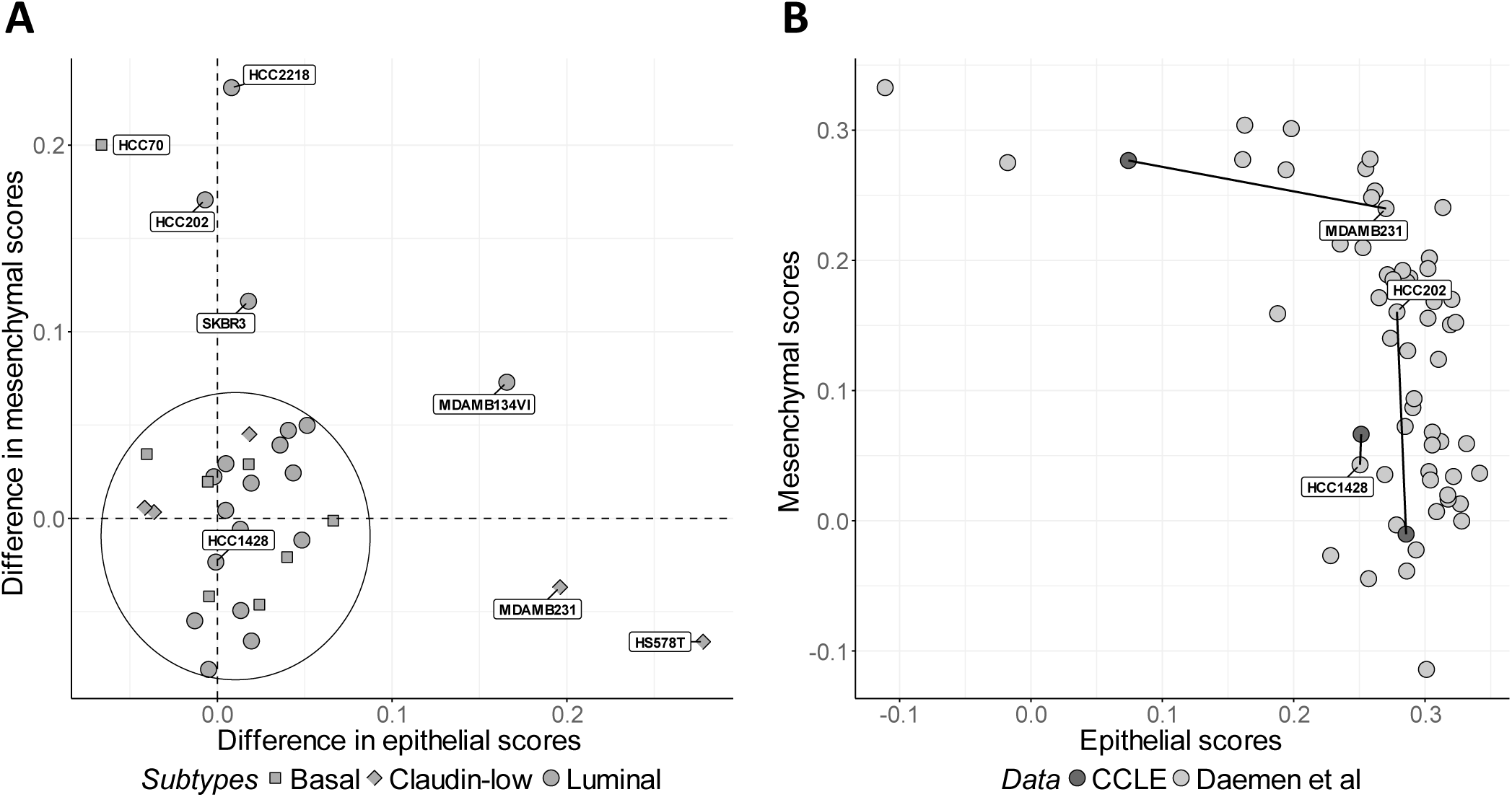
(A) Differences in epithelial and mesenchymal sores for 32 overlapping breast cancer cell lines between Daemen et al and the CCLE datasets. The majority of the cell lines show relatively consistent scores in these two data sets (circled in the lower left corner). (B) The HCC1428 cell line has very similar scores in each dataset, while the MDAMB231 cell line has a large shift in epithelial score, and the HCC202 cell line has a large shift in mesenchymal score.

### Application of simple scoring method

#### Obtaining landscapes of molecular phenotypes

Scoring samples against multiple molecular signatures and plotting them in 2D or 3D can be useful to stratify samples based upon the associated molecular phenotypes of samples. For example, scoring the TCGA breast cancer samples (*n_Samples_* = 1091 RNA-seq), and breast cancer cell lines from A Daemen, OL Griffith, LM Heiser, NJ Wang, OM Enache, Z Sanborn, F Pepin, S Durinck, JE Korkola, M Griffith, et al. [17] (*n_Samples_* = 64 RNA-seq) against their respective mesenchymal and epithelial signatures from TZ Tan, QH Miow, Y Miki, T Noda, S Mori, RY Huang and JP Thiery [24], provides more refined stratification of patients and cell lines. For instance, only a subset of the aggressive claudin-low breast cancer subtype consistently shows high mesenchymal and low epithelial scores across the independent data sets (Figure 3) [28]. These samples have different expression patterns compared to samples with high epithelial and low mesenchymal scores, representing a subset of less aggressive subtypes (Luminal). Each of these sub-groups can be further analysed, for example by comparing different omics data between these sub-groups, or by examining their associations with pre-clinical outcomes such as drug response.

**Figure 3.**
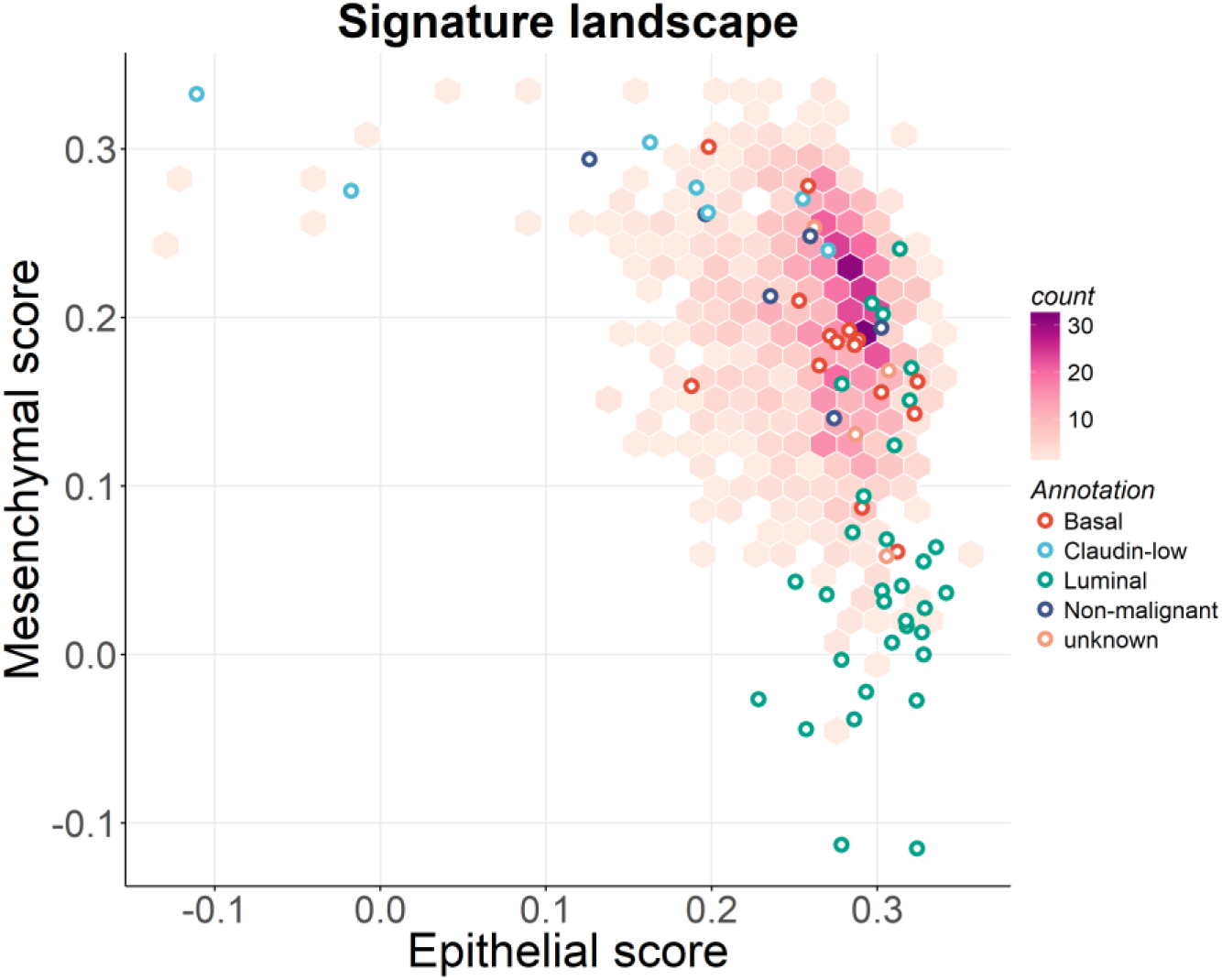
Epithelial and mesenchymal scores obtained from *singscore* for the TCGA breast cancer samples (*hexbin density plot*) and a collection of breast cancer cell lines (*circle markers*, *coloured by subtype*). Note that as per the original study by Tan *et al*., the epithelial and mesenchymal signatures are distinct (but overlapping) for tumours and cell lines.

#### Assessment of scores: beyond a single value

When ranking gene expression values of a sample in increasing order, a higher mean rank of the expected up-regulated gene set represents a higher concordance of the sample to the gene set, while lower ranks of expected down-regulated genes are associated with a higher concordance of the sample to the gene set. Considering where gene ranks would sit within a sample with the maximum theoretical scores (see *Methods*), the expected up- and down-regulated gene sets should form an approximately bimodal distribution, with higher ranks for the up- and lower ranks for the down-set (Figure 4A, sample 3). For samples that are not strongly concordant with a signature, the distributions of genes are less coordinated, and often relatively uniform (Figure 4A, sample 2).

**Figure 4.**
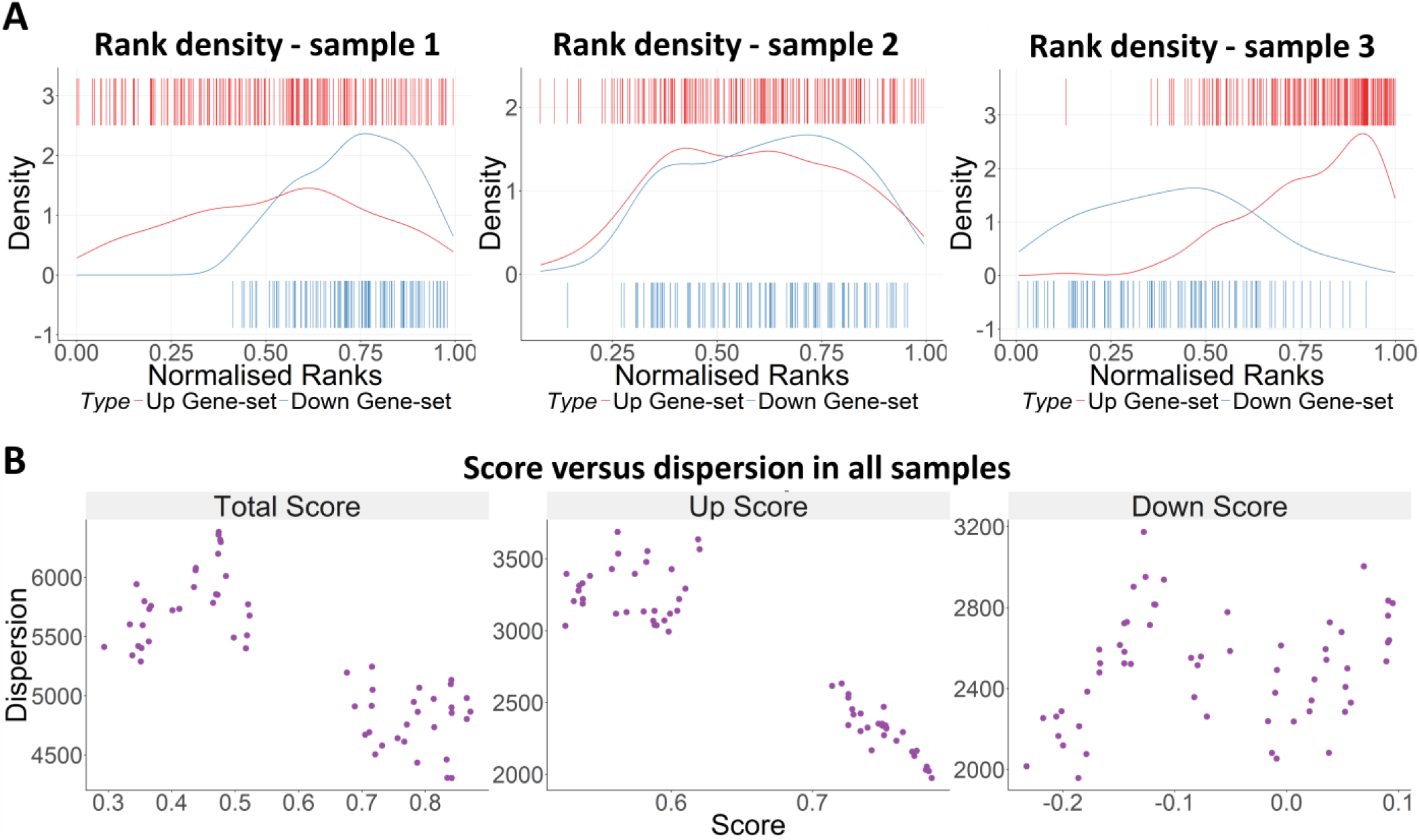
(A) Three microarray samples from the TGFβ-EMT data set [1] with low, medium and high scores for the TGFβ-EMT signature. (B) Scatter plots demonstrating the relationship between rank dispersions (MAD) and scores (obtained by *singscore)* for the total scores (combined up- and down-set scores), up-set scores and down-set scores.

To easily visualise the rank distribution of genes in the up- and down-gene sets, the *singscore* package provides static and interactive plots that display density and barcode plots for gene ranks in individual samples (Figure 4A). These plots help to interpret the score in the context of the ranked genes, and often demonstrate that up- and down-gene sets can vary in their dispersion, contributing to the range of ranks we observe. We often see that a low scoring sample may have an inverted pattern of expression (Figure 4A, sample 1), or have no concordance with the gene set at all (*i.e*. randomly distributed genes across the ranks; Figure 4A, sample 2).

To illustrate these differences, we calculate median absolute deviation (MAD) of the gene set ranks to estimate relative rank dispersion. Plotting scores against dispersion for the samples in the TGFβ-EMT data shows that samples with a high total score also have lower dispersion, demonstrating more coordinated changes in the up- and/or down-sets in these samples (Figure 4B). It is also possible to look at the rank dispersion of the up- and down-sets to assess the performance of each of these sets separately. Figure 4B shows that genes in the up-set of TGFβ-EMT signature are more coordinated in samples compared to the down-set.

## Discussion

We have described a rank-based single sample gene set scoring method, implemented in the *singscore* package. Our method can easily be applied on any high throughput transcriptional data from microarray or RNA-seq experiments. While our method is non-parametric, it needs read counts to be adjusted for gene lengths and ideally, GC content biases, because these may alter gene ranks within individual samples. Accordingly, RNA abundance data formatted as RPKM, TPM, or RSEM can be used, with or without log-transformation, and/or filtering for genes with very low expression.

Using microarray and RNA-seq platforms of the TCGA breast cancer data, we show that our *singscore* approach yields stable scores for individual samples because they are treated independently from other samples, in contrast to GSVA, PLAGE, z-score, and ssGSEA. Although modifying ssGSEA to exclude the final normalization step also shows high stability, the lack of a single sample normalization/scaling step in ssGSEA_!Norm_ hinders interpretability of scores, that is, it is difficult to make a distinction between a moderate score and a high score. Additionally, even with modification, ssGSEA cannot actually be applied to a single sample. In contrast, *singscore* includes per sample normalisation and scaling by considering the theoretical minimum and maximums for scores in each sample, and can be applied to a single sample in isolation.

We show that current implementations of both ssGSEA_!Norm_ and ssGSEA through the GSVA package are computationally much slower than all the other methods for scoring samples against a large number of random signatures (Suppl. Figure 1), while our approach is fast. The poor performance of PLAGE (Suppl. Figure 3) for signatures with both up- and down-sets, even for the large number of samples, may be due the fact that this method is a “pathway” analysis approach in which the direction of changes in pathway genes are not of interest. We noticed this with correlation of PLAGE scores being high but negative compared to other scores for the epithelial signature. Although we have not assessed this, the performance of this method may increase if the up- and down-sets are provided as one gene set.

We compared breast cancer cell lines overlapping between the CCLE and Daemen et al data and showed high consistency in epithelial and mesenchymal scores obtained by our scoring approach for the majority of cell lines (Figure 2A). Because only a small subset of cell lines shows relatively larger differences in epithelial or mesenchymal scores between the two data sets it is tempting to speculate that variations in scores are not due to the differences in technical or computational pipelines, which would have affected all the cell lines in the analysis. Rather it is possible that differences in scores reflects real biological variation in the molecular phenotypes of some cells: the more-variable cell lines within the Daemen et al. data have hybrid epithelial-mesenchymal phenotypes (*i.e*. high epithelial and mesenchymal scores); these cell lines may be more dynamic in terms of epithelial-mesenchymal plasticity, which may explain the variability in their EMT phenotype under different experimental conditions. Interestingly, all the cell lines with relatively large variations in mesenchymal axis with smaller differences in epithelial scores are luminal cell lines which in general are shown to have strong epithelial phenotype (Figure 3, and [28]), while most cell lines with large shifts on the epithelial axis and little change in mesenchymal scores are claudin low cell lines which have been shown to be strongly mesenchymal (Figure 3, and [28]).

It is also important to note that there are a few more recent single sample analysis approaches, including multiple omics gene set analysis (moGSA) [29] and personalized pathway alteration analysis (PerPAS) [30], which have not been discussed here, as they are fundamentally different from our approach. For example, moGSA needs multiple types of large-scale (‘omics’) data for each individual sample and it integrates information across these data using multivariate factor analysis [29], and thus it may not be applicable for samples with only transcriptomic data. Alternately, PerPAS needs topological information for each gene to perform pathway analysis, and further uses either a control sample or a cohort of samples based on which the gene expression data in single sample are normalised [30]; these requirements make this method unsuitable for many available datasets.

Methods that need a large number of samples to calculate a precise and stable score for each individual sample often need to be re-run across a large data set whenever a few new samples are required to be scored. It adds an extra layer of complexity to the already complicated scoring methods. Our single sample scoring method provides a simple and easy-to-understand pipeline which performs as well as the other comparable scoring methods in large data sets, and outperforms other methods in smaller data sets. Several visualization options at both bulk and single sample level are provided to enable users to explore genes, gene signatures, and samples in more depth.

## Conclusion

In the context of personalised medicine, we often have data from an individual patient, or from a small number of samples in pre-clinical experiments. Current scoring methods are parametric and/or depend on large number of samples to produce stable scores, while *singscore* generates scores that are stable across a range of sample sizes and numbers of measured genes. This is due to it being a non-parametric, rank-based, and truly single sample method. Moreover, scores generated by our method are less likely to be influenced by batch effects across samples and can be normalised at a single sample level, resulting in easily-interpretable scores.

## Acknowledgements

Results shown are in part based upon data generated by the Cancer Cell Line Encyclopedia (http://www.broadinstitute.org/ccle) and TCGA Research Network: (http://cancergenome.nih.gov/).The authors would like to thank Prof. Terry Speed for his advice and support.

## Declaration

### Ethics approval and consent to participate

NA

### Consent for publication

NA

### Availability of data and material

An *R/Bioconductor* package *singscore* is under development and the current version can be downloaded from: https://github.com/DavisLaboratory/singscore

### Authors’ contributions

- **Conception and design:** M. Foroutan, D. Bhuva, K. Horan, J. Cursons, M.J. Davis
- **Development of methodology:** M. Foroutan, D. Bhuva, K. Horan, J. Cursons, M.J. Davis **Analysis and interpretation of data (e.g., statistical analysis, biostatistics, computational analysis):** M. Foroutan, D. Bhuva, K. Horan, R. Lyu, J. Cursons, M.J. Davis
- **Writing, review, and/or revision of the manuscript:** M. Foroutan, D. Bhuva, K. Horan, J. Cursons, M.J. Davis
- **R package development:** R. Lyu, M. Foroutan, D. Bhuva, K. Horan
- **Study supervision:** J. Cursons, M.J. Davis

## List of abbreviations

TGFβ: transforming growth factor beta
EMT: epithelial-mesenchymal transition
ssGSEA: single sample gene set enrichment analysis
GSVA: gene set variation analysis
PLAGE: pathway-level activity of gene expression
moGSA: multiple omics gene set analysis
PerPAS: personalized pathway alteration analysis
CCLE: cancer cell line encyclopaedia

## References

1. Foroutan M, Cursons J, Hediyeh-Zadeh S, Thompson EW, Davis MJ: A Transcriptional Program for Detecting TGFβ-Induced EMT in Cancer. Molecular Cancer Research 2017, 15(5):619–631.

2. Barbie DA, Tamayo P, Boehm JS, Kim SY, Moody SE, Dunn IF, Schinzel AC, Sandy P, Meylan E, Scholl C et al: Systematic RNA interference reveals that oncogenic KRAS-driven cancers require TBK1. Nature 2009, 462(7269):108–112.

3. Hanzelmann S, Castelo R, Guinney J: GSVA: gene set variation analysis for microarray and RNA-seq data. BMC Bioinformatics 2013, 14:7.

4. Tomfohr J, Lu J, Kepler TB: Pathway level analysis of gene expression using singular value decomposition. BMCBioinformatics 2005, 6:225.

5. Lee E, Chuang HY, Kim JW, Ideker T, Lee D: Inferring pathway activity toward precise disease classification. PLoS Comput Biol 2008, 4(11):e1000217.

6. Oshlack A, Wakefield MJ: Transcript length bias in RNA-seq data confounds systems biology. Biol Direct 2009, 4:14.

7. Zheng W, Chung LM, Zhao H: Bias detection and correction in RNA-Sequencing data. BMC Bioinformatics 2011, 12:290.

8. Benjamini Y, Speed TP: Summarizing and correcting the GC content bias in high-throughput sequencing. Nucleic Acids Res 2012, 40(10):e72.

9. Patro R, Duggal G, Love MI, Irizarry RA, Kingsford C: Salmon provides fast and bias-aware quantification of transcript expression. Nat Methods 2017, 14(4):417–419.

10. Bray NL, Pimentel H, Melsted P, Pachter L: Near-optimal probabilistic RNA-seq quantification. Nat Biotechnol 2016, 34(5):525–527.

11. Roberts A, Trapnell C, Donaghey J, Rinn JL, Pachter L: Improving RNA-Seq expression estimates by correcting for fragment bias. Genome Biol 2011, 12(3):R22.

12. Wickham H: ggplot2: Elegant Graphics for Data Analysis. 2009.

13. Sievert C, Parmer C, Hocking T, Chamberlain S, Ram K, Corvellec M, Despouy P: plotly: Create Interactive Web Graphics via “plotly.js”. In.; 2017.

14. Hänzelmann S, Castelo R, Guinney J: {GSVA}: gene set variation analysis for microarray and {RNA-Seq} data. BMC Bioinformatics 2013, 14:7.

15. Cancer Genome Atlas N: Comprehensive molecular portraits of human breast tumours. Nature 2012, 490(7418):61–70.

16. Barretina J, Caponigro G, Stransky N, Venkatesan K, Margolin AA, Kim S, Wilson CJ, Lehar J, Kryukov GV, Sonkin D et al: The Cancer Cell Line Encyclopedia enables predictive modelling of anticancer drug sensitivity. Nature 2012, 483(7391):603–607.

17. Daemen A, Griffith OL, Heiser LM, Wang NJ, Enache OM, Sanborn Z, Pepin F, Durinck S, Korkola JE, Griffith M et al: Modeling precision treatment of breast cancer. Genome Biol 2013, 14(10):R110.

18. UCSC Cancer Genome Browser [https://genome-cancer.ucsc.edu]

19. Cancer Cell Line Data Repository [https://www.synapse.org/#!Synapse:syn5612998]

20. Gene Expression Omnibus [https://www.ncbi.nlm.nih.gov/geo/]

21. NCBI SRA Toolkit [https://www.ncbi.nlm.nih.gov/sra/docs/toolkitsoft]

22. Liao Y, Smyth GK, Shi W: The Subread aligner: fast, accurate and scalable read mapping by seed-and-vote. Nucleic Acids Research 2013, 41:e108.

23. Robinson MD, McCarthy DJ, Smyth GK: edgeR: a Bioconductor package for differential expression analysis of digital gene expression data. Bioinformatics 2010, 26:-1.

24. Tan TZ, Miow QH, Miki Y, Noda T, Mori S, Huang RY, Thiery JP: Epithelial-mesenchymal transition spectrum quantification and its efficacy in deciphering survival and drug responses of cancer patients. EMBO Mol Med 2014, 6(10):1279–1293.

25. Liberzon A, Subramanian A, Pinchback R, Thorvaldsdóttir H, Tamayo P, Mesirov JP: Molecular signatures database (MSigDB) 3.0. Bioinformatics 2011, 27(12):1739–1740.

26. Subramanian A, Tamayo P, Mootha VK, Mukherjee S, Ebert BL, Gillette MA, Paulovich A, Pomeroy SL, Golub TR, Lander ES: Gene set enrichment analysis: a knowledge-based approach for interpreting genome-wide expression profiles. Proceedings of the National Academy of Sciences 2005, 102(43):15545–15550.

27. Harell F: Regression modeling strategies. Guildford (UK): Springer 2001.

28. Cursons J, Pillman KA, Scheer K, Gregory PA, Foroutan M, Zadeh SH, Toubia J, Crampin EJ, Goodall GJ, Bracken CP: Post-Transcriptional Control Of EMT Is Coordinated Through Combinatorial Targeting By Multiple microRNAs. bioRxiv 2017:138024.

29. Meng C, Kuster B, Peters B, Culhane AC, Gholami AM: moGSA: integrative single sample gene-set analysis of multiple omics data. bioRxiv 2016:046904.

30. Liu C, Lehtonen R, Hautaniemi S: PerPAS: Topology-Based Single Sample Pathway Analysis Method. IEEE/ACM Transactions on Computational Biology and Bioinformatics 2017.

